# Action representation in the mouse parieto-frontal network

**DOI:** 10.1101/646414

**Authors:** Tuce Tombaz, Benjamin A. Dunn, Karoline Hovde, Ryan J. Cubero, Bartul Mimica, Pranav Mamidanna, Yasser Roudi, Jonathan R. Whitlock

## Abstract

The posterior parietal cortex (PPC), along with anatomically linked frontal areas, form a cortical network which mediates several functions that support goal-directed behavior, including sensorimotor transformations and decision making. In primates, this network also links performed and observed actions via mirror neurons, which fire both when an individual performs an action and when they observe the same action performed by a conspecific. Mirror neurons are thought to be important for social learning and imitation, but it is not known whether mirror-like neurons occur in similar networks in other species that can learn socially, such as rodents. We therefore imaged Ca^2+^ responses in large neural ensembles in PPC and secondary motor cortex (M2) while mice performed and observed several actions in pellet reaching and wheel running tasks. In all animals, we found spatially overlapping neural ensembles in PPC and M2 that robustly encoded a variety of naturalistic behaviors, and that subsets of cells could stably encode multiple actions. However, neural responses to the same set of observed actions were absent in both brain areas, and across animals. Statistical modeling analyses also showed that performed actions, especially those that were task-specific, outperformed observed actions in predicting neural responses. Overall, these findings show that performed and observed actions do not drive the same cells in the parieto-frontal network in mice, and suggest that sensorimotor mirroring in the mammalian cortex may have evolved more recently, and only in certain species.

## Introduction

A key function of any motor system is the rapid and flexible production of actions in response to external stimuli, including the behavior of other individuals. Having robust representations of performed and observed behaviors has therefore been hypothesized to add survival value in a number of species since it facilitates an array of behavioral functions, including optimal action selection, gaining access to food sources or avoiding predators (1). However, which neural circuits integrate performed and observed actions, and how, are not well understood. In different species of primates and songbirds, a striking manifestation of such interactions has been described in the form of mirror neurons. Mirror neurons, first characterized in pre-motor cortex (2, 3) then PPC (4) in monkeys, and later reported in humans (5) and birds (6), respond reliably both when an individual performs a specific action and when they observe the same action performed by a conspecific. Based on these properties, they have been postulated to enable skills requiring conjoint coding of observed and performed behaviors, such as imitation and action understanding (7, 8). The striking specificity of mirror coding requires that sensory and motor processing streams are combined precisely at the level of single cells, which prompts the question as to how such a mechanism arose originally. That is, did prototypical sensory and motor processing pathways become linked early in evolution, in which case most species should exhibit sensorimotor mirror matching, or is it a more recent adaptation suited to the needs of a few specific niches?

To address this question, we tested whether neurons in PPC and frontal motor cortex (M2) of mice encode the performance and observation of unrestrained motor behaviors. We chose rodents since they fall between primates and avians phylogenetically, they can socially acquire both sensorimotor and fear-based behaviors (9–14), and they have proven effective models for studying the neurobiology of empathetic social learning (15–19). Emerging evidence also suggests that PPC and M2 in rodents, like primates, comprise a cortical network supporting several aspects of goal-directed behavior, including decision making (20, 21), sensorimotor transformations (22, 23), and movement planning (24, 25). Rodent models also bring methodological advantages, including large-scale neural recordings in unrestrained subjects, which enables the analysis of neural ensemble dynamics during any number of self-initiated actions. In turn, it is possible to uncover intrinsic features of neural population activity driven by behavior, such as state space structure (26), independently of experimenter bias.

Here, we used miniaturized, head-mounted fluorescent microscopes (27) to image the activity of hundreds of individual neurons at a time while mice performed or observed pellet reaching and wheel-running tasks. First, using dimensionality reduction (28), we saw clear differences in the structure of ensemble responses during performed and observed behaviors. This motivated the subsequent quantification of single-cell selectivity to specific behaviors using shuffling analyses as well as statistical modeling with a generalized linear model (GLM). All tests indicated that PPC and M2 were driven strongly by performed behaviors, similar to what has been shown in more stereotypical tasks (26), but extended here to freely behaving animals. The neural coding of observed behavior, on the other hand, was below chance levels in both brain areas, even in neurons with strong performance correlates. These results indicate that the representation of the observed actions we tested occurs outside the parieto-frontal circuit in mice, which suggests a divergence in action recognition mechanisms between primates and rodents. By extension, this supports the view that sensorimotor mirroring evolved independently in birds and primates.

## Results

To determine whether neurons in the mouse PPC and M2 reliably responded to the performance and observation of the same set of behaviors, we used one-photon epifluorescence microscopy to image the activity of neuronal ensembles expressing the genetically encoded calcium indicator GCaMP6m (AAV1.Syn.GCaMP6m.WPRE.SV40) via AAV-mediated transfection (921 neurons in PPC in 4 mice; 852 neurons in M2 in 4 mice; Fig. S1, Table 1). Cellular responses were monitored through a chronically implanted gradient refractive index lens attached to a prism (Fig. 1 *A* and *C*). All animals were trained to perform the pellet reaching task (29) in an 8.5 x 15 x 20 cm box (Fig. 1*B*), in which they were taught to reach through a 1 cm diameter hole to grasp food pellets (Fig. 1*B*). They were trained to asymptotic performance levels prior to experimental recordings (maximum of 10 days; Methods) and, concurrently, were habituated to head-fixation and to observe a sibling perform the same task. In the experiments, each animal’s cortical activity was imaged during four sessions, with performance (P) and observation (O) conditions interleaved (following a P1-O1-P2-O2 scheme). In parallel, we recorded from each mouse while they behaved freely in a wall-less open arena (30 x 30 cm) with a running wheel, and while they observed a sibling doing the same (Fig. 1*B*). The calcium imaging data were paired with high-resolution behavioral recordings made during both performance and observation sessions.

**Fig. 1.**
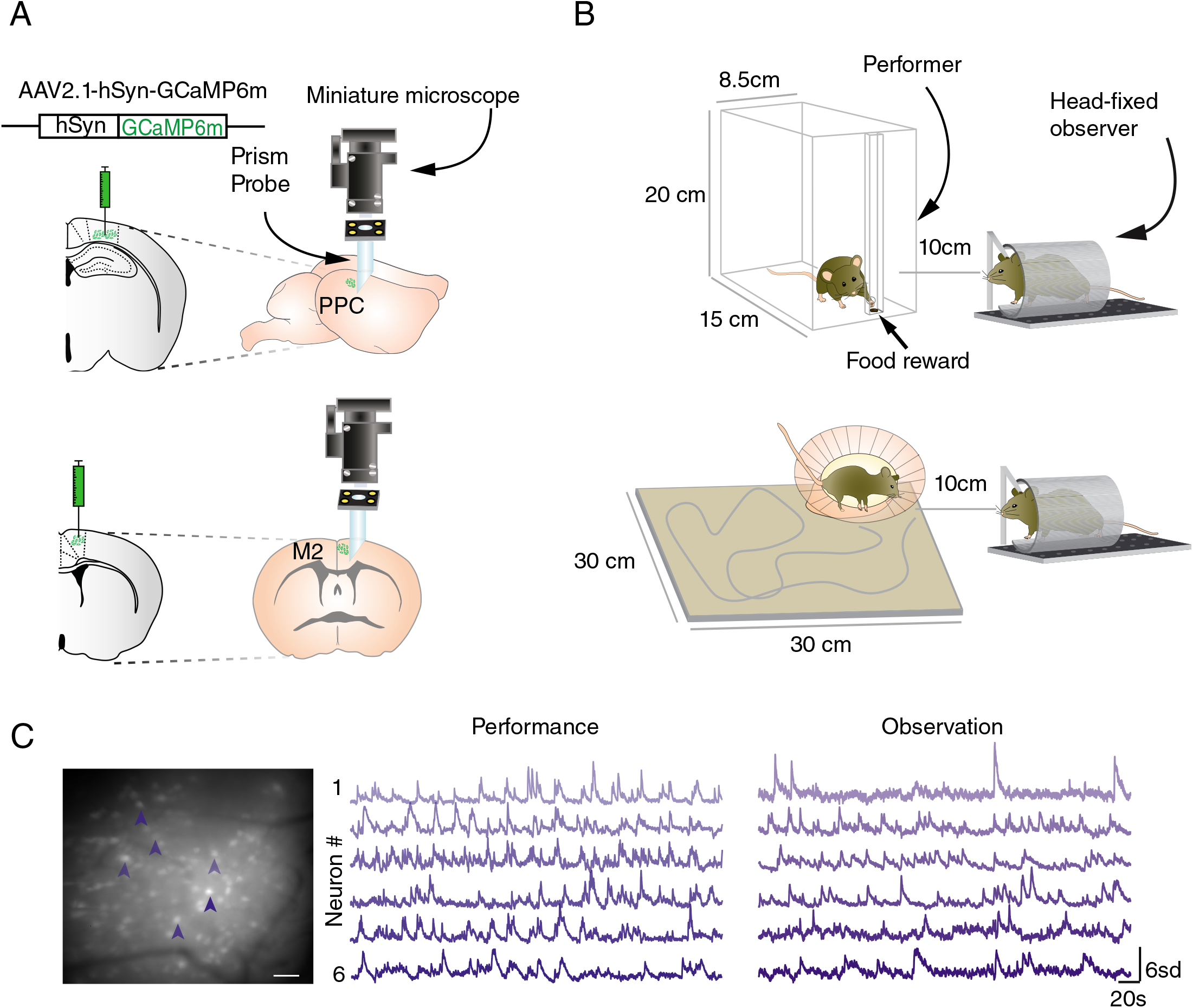
Experimental paradigm for imaging neural populations in PPC and M2 in freely behaving mice. (*A*) PPC and M2 were transfected virally to express GCaMP6m (*Left*), and miniature prism probes were implanted to image tangentially across cortical layers (*Right*) during different behavioral tasks. (*B*) In the experiments, mice alternated between performing and observing a conspecific in a pellet-reaching task (*Top*) and wheel-running task in an open arena (*Below*). Dynamic calcium fluctuations were monitored in each mouse during four 10-min recording sessions, two of which were during performance and two during observation of each task. (*C*) (*Left*) Average of 500 images of the entire FOV after image pre-processing. Scale bar, 100μm. Shaded arrows indicate 6 cells whose calcium traces are shown (*Middle*) during performance and (*Right*) observation of the pellet-reaching task.

Having imaged large ensembles of neurons in PPC and M2, as a prelude to our analysis, we visualized how performance and observation conditions affected the population activity. To this end, we applied the uniform manifold approximation and projection (UMAP) method on downsampled population activity vectors (Methods) (28). As shown in Fig. 2 *A* and *B*, this revealed structural discrepancies in the dimensionally-reduced activity space between performance and observation sessions, with population activity states being closer to each other for time points belonging to the same behavior during performance, but not observation conditions (Movie S1). We measured the degree to which time points labeled by the same behaviors were clustered using the Dunn Index (Methods), which produced clustering indices between 2.4 and 10.9 times higher during performance than observation sessions across animals (3 mice in PPC, 1 mouse in M2). This suggested that there were clear signatures of the representation of performed behavior but not observed behavior in PPC and M2 activity. Due to the dependence of the quantitative aspects of the UMAP results on several initial parameters, such as the dimension of the projective space, a more careful quantification of these effect required going beyond this visualization, which is what we report in the rest of the paper.

**Fig. 2.**
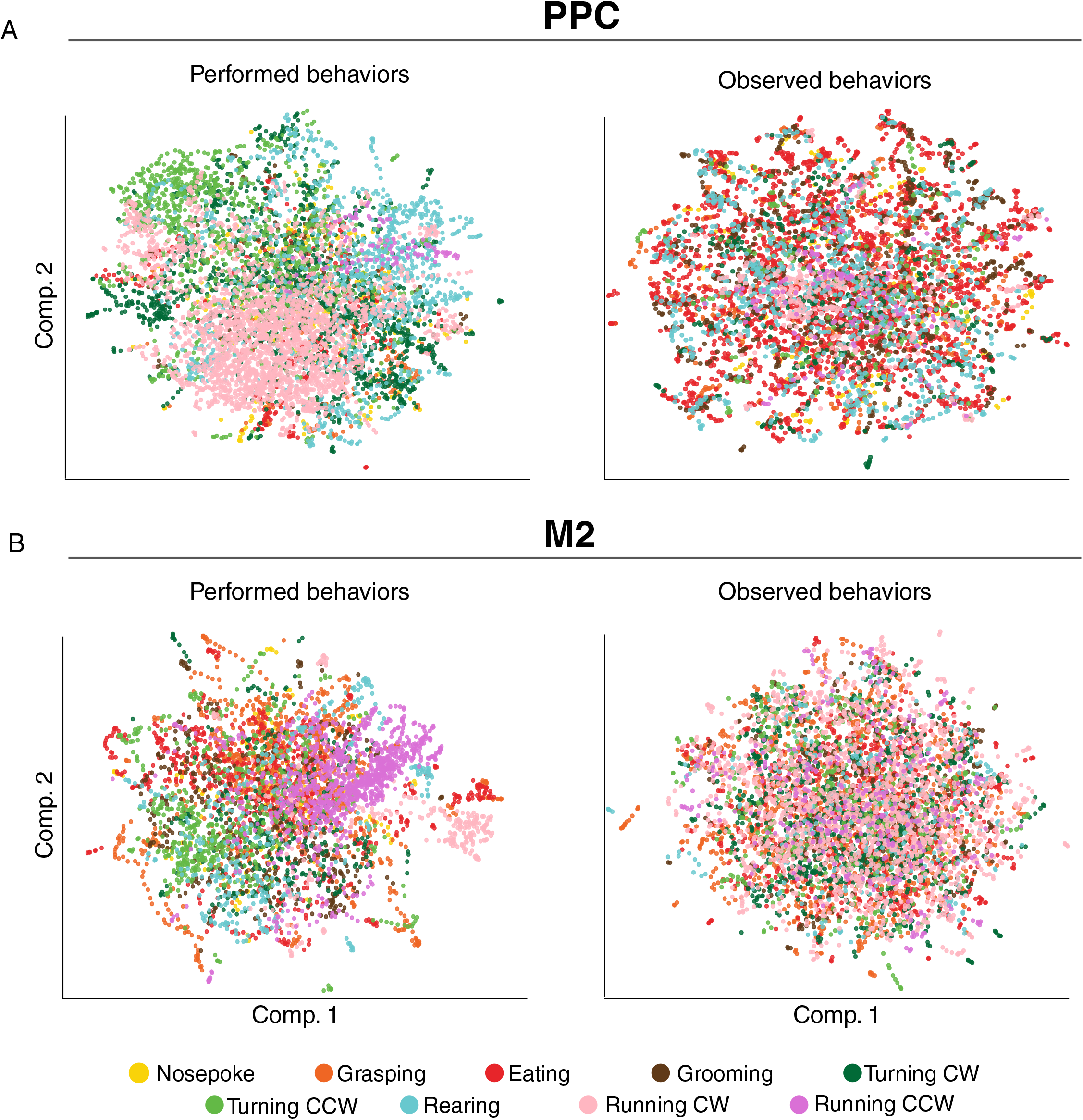
UMAP projections of population activity in both PPC and M2 reveal structural segregation for performed but not observed behaviors. (*A*) PPC ensemble activity separated in the reduced dimensional space during specific performed behaviors, including wheel-running (beige dots), counter-clockwise turning (light green) and rearing (blue). By contrast, the distribution of points during observed behaviors (*Right*) was spread homogenously in UMAP space. Each dot corresponds to the activity state of the entire population of recorded neurons at a given time point; color-coding for each behavior is shown at bottom. (*B*) Recordings from M2 were similar to *A*, showing a stronger tendency to cluster during performed than observed behaviors.

To determine if the UMAP results reflected behavioral selectivity at the single-cell level, we quantified the tuning of individual neurons to different actions the animals engaged in while performing the tasks. We labelled the onset and offset of discrete, recurring behaviors, including turning left or right, nose poking, grasping to eat, eating, rearing or grooming (Fig. 3 *A*; Movie S2; Methods). A cell was considered stably tuned to a behavior if its in-behavior event rate exceeded 95% of the shuffled in-behavior rates in two consecutive performing sessions (Methods). Approximately half the neurons in both PPC (430 of 921 cells; 46.6% in 4 mice) and M2 (439 of 852 cells, 51.5%, 4 mice) were reliably driven by performed behaviors (Fig. 3 *B* and *C*; Table S1). While the majority of neurons were uniquely tuned to individual behaviors, subsets of cells were selective for multiple actions, and in all cases, tuning was invariant to the duration of the behavior (Fig. S2). In the open field task, 67 of 724 PPC cells (9.3%; 3 mice) in PPC were stably tuned to clockwise (CW) and counter-clockwise (CCW) wheel running, while 21 out of 216 neurons (9.7%, running CCW only; 1 mouse) were stably tuned in M2. The proportion of cells representing each behavior varied between animals, with larger groups of cells encoding turning in both PPC and M2, and a larger proportion of cells tuned to grasping in M2 than PPC (Figs. 3 *B-D*, S2).

**Fig. 3.**
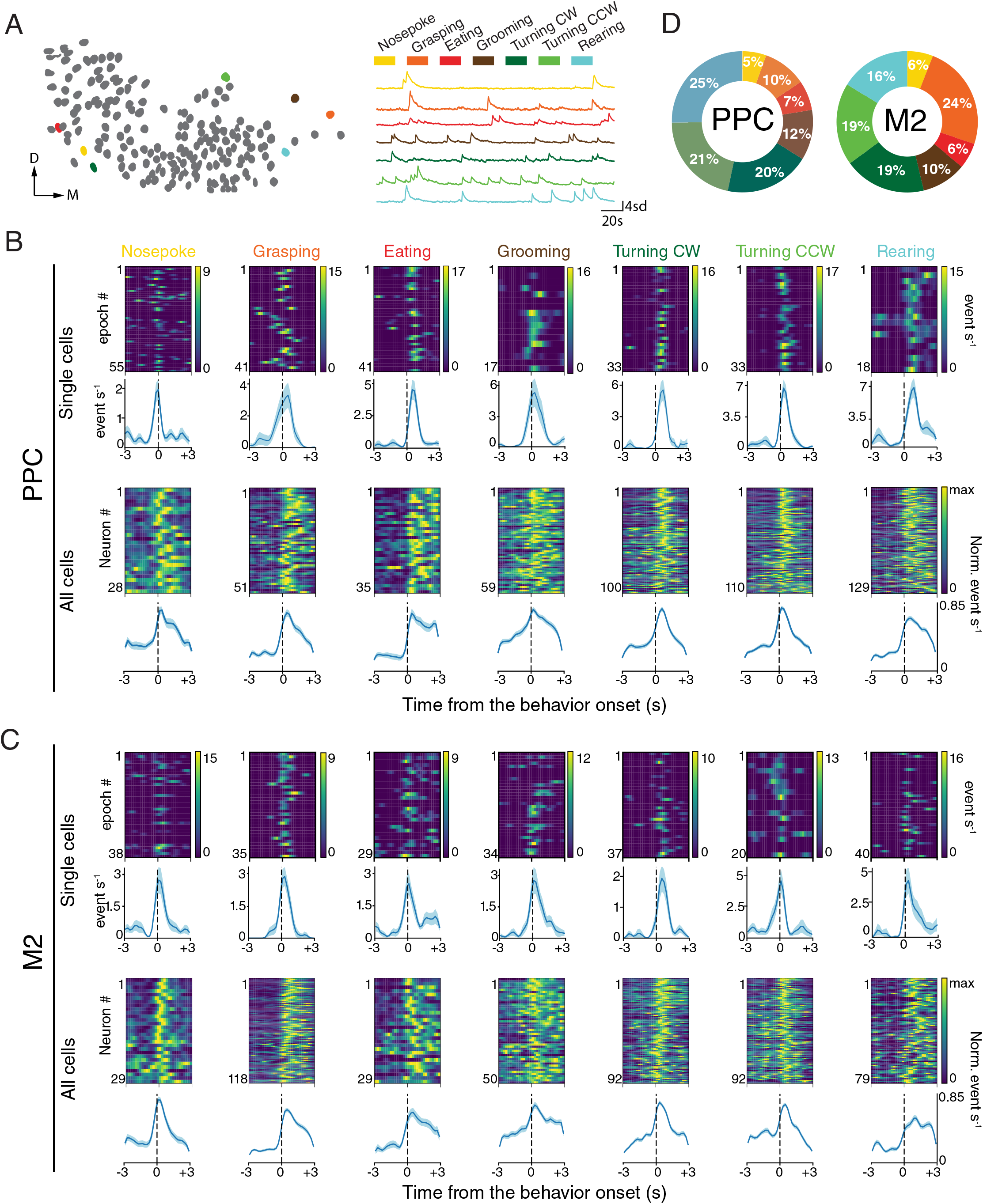
Cell populations in PPC and M2 robustly encode actions performed in the pellet-reaching task. (*A*) Representative neural map (*Left*) and Ca^2+^ transients of 7 PPC neurons (*Right*) tuned to each of the behaviors in the pellet reaching task; color coding for each behavior is shown above. (*B*) (*Top*) Temporal profiles of behaviorally evoked responses of single cells for each behavior are shown as heat maps; immediately beneath are behaviorally aligned average activity rates for each cell over the entire session. (*Bottom*) Normalized responses for all behaviorally tuned PPC neurons from all animals aligned to behavior onset; population means are shown in the row underneath. Color bars indicate max event/s; blue shaded regions around averaged rates denote ± SEM. (*C*) Same as *B*, for single cells (*Top row*) and cell populations (*Bottom row*) imaged in M2. (*D*) Color coded pie charts indicating the proportion of stably tuned neurons for each behavior; a total of 1674 cells were imaged in 4 mice in PPC; 1082 cells were imaged in 4 mice in M2.

The heterogeneity of tuning properties, and the tuning of some cells to multiple behaviors, raised the question as to whether cells with similar coding clustered anatomically, as suggested by prior work in parietal and motor areas in different mammalian species (30–32). Ensemble imaging allowed us to assess the spatial micro-organization of behaviorally responsive neurons according to their tuning preference in each brain region of each animal. However, an analysis of the quality of clustering by behavioral tuning (Dunn Index; Methods) showed no clear tendency of grouping between cells with similar properties, nor any clear mapping based on cortical depth or location in the imaging field of view (Fig. S3 and S4).

Since PPC and M2 showed robust tuning to a variety of performed behaviors, we next assessed whether they responded during observation of the same actions. We compared trial-averaged responses to specific behaviors across all four recording sessions: P1, O1, P2 and O2 (Fig. 4 *A* and *B* upper panels). However, in both brain areas and across mice, we saw negligible neural tuning to observed actions, irrespective of whether the cells stably encoded performed actions (Fig. 4 *A* and *B* lower panels, Fig.S5). Specifically, 15 of 921 neurons (1.6%) in PPC and 13 out of 852 neurons (1.5%) in M2 exhibited stable observational correlates for the pellet reaching task, even though the total amount of time the animals spent observing behaviors was comparable to the time spent performing them (Table S1). To test whether the proportion of cells reliably tuned to observed behaviors exceeded chance levels, we paired neural activity with behavior labels from the wrong sessions and computed false positive rates in this manner for all sessions and all animals (Methods). This approach identified 27/921 (2.9%) PPC cells and 32/852 (3.8%) M2 cells as stably tuned to mismatched observed behaviors, demonstrating that the low number of stable observational correlates was less than expected by chance (PPC: U= 286.5, p > 0.05, M2: U = 228, p > 0.05; Mann-Whitney U test). Similarly, in the open field task, only three out of 724 neurons (0.4%) and one out of 216 neurons (0.5%) had reliable observational tuning to running behaviors in PPC and M2, respectively. Fewer than 1% of cells had stable, matched correlates for performed and observed actions in the pellet reaching task in either area, which again was below mismatched data rates (U = 364, p > 0.05 for PPC; U = 287.5, p > 0.05 for M2). Moreover, no cells showed matched tuning for wheel-running behavior in the open field task (Table S1).

**Fig. 4.**
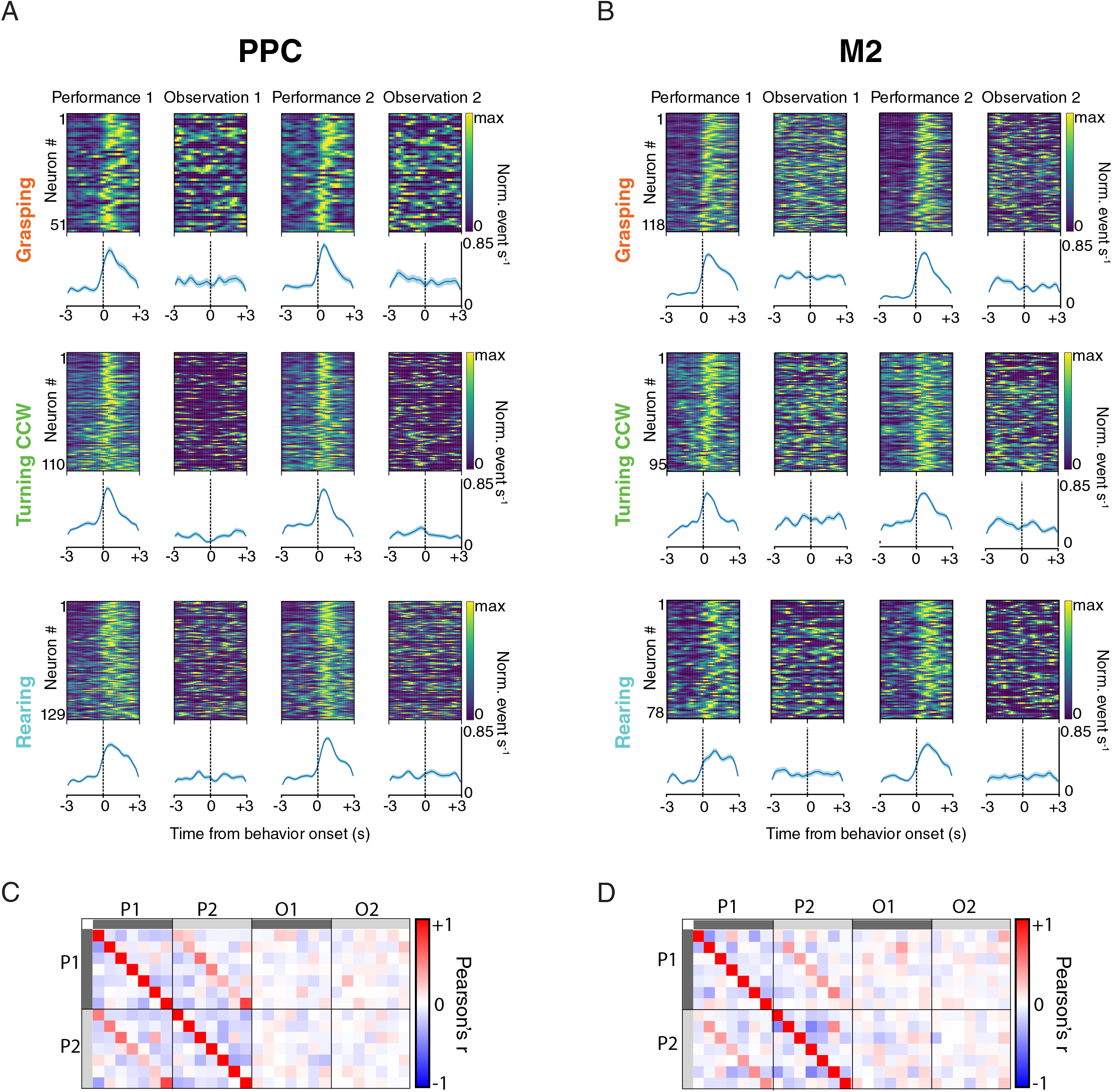
Neural ensembles in PPC and M2 stably represent performed, but not observed actions. (*A*) (*Above*) Session-averaged Ca^2+^ responses of individual cells aligned to the onset of specific actions in the pellet- reaching task, and ranked by z-scored firing rate during the first performance session (P1). (*Below*) Population average (± SEM) of responses of all cells for each behavior. Virtually none of the cells with stable correlates across the two performance sessions responded when the same actions were observed (Observation sessions 1 and 2), yielding a flat activity rate in the ensemble average. (*B*) Same as *A*, but for cells recorded in M2. (*C*) Correlation matrices, with each square corresponding to a particular behavior, show the sustained specificity of behavioral tuning in PPC across performance sessions (P1 and P2). The conserved correlation structure is reflected by the red diagonal in P1 vs. P2, which is notably absent across performance and observation conditions. (*D*) Same as *C*, for recordings in M2.

To investigate whether the lack of neural responsiveness to observed actions stemmed from fluctuations in arousal state, we measured variations in pupil diameter, a proxy for arousal and attention (33), in a subset of mice. Since prior work established that contraction of the pupil is associated with reduced attentiveness and neural responsiveness (34), we restricted our analyses of observation sessions to exclude epochs when the pupil diameter was smallest (Fig. S6; n = 3 animals). Consistent with our prior findings, however, this did not affect the number of cells showing stable tuning (9 of 621 cells (1.5%) with all timepoints included, 8 cells (1.3%) when excluding pupil contraction), indicating the lack of effect did not relate to low arousal of the observers.

To further assess whether activity patterns during performance sessions related to observation, we sought to characterize how well cellular activity could be predicted from one task condition to another. When cells were selected based on their behavioral tuning in the first performing session, and their z-scored firing rates were correlated to those in the second performing session, we saw in every case that the responses of cells correlated positively (Pearson’s correlations for same-behavior comparisons ranged from 0.22 to 0.75 for PPC and 0.08 to 0.48 for M2; Fig. 4*C*). Likewise, selecting cells based on their tuning in the second session and correlating those rates back to the first yielded similar results (r-values ranged from 0.20 to 0.71 in PPC and 0.1 to 0.44 for M2; Fig. 4*C*). By comparison, the correlations of activity rates between performance and observation sessions centered around zero in all cases (r-values ranged from −0.09 to 0.12 for PPC and −0.24 to 0.22 for M2; Fig. 4*C*).

Lastly, we wished to determine the extent to which each of the behaviors explained the activity rates of the cells during performance and observation conditions, for which we used a generalized linear model (GLM) framework (Methods) (35). The model was designed to incorporate all labelled behaviors as predictors of each neuron’s time-varying activity. To quantify how well the behavioral variables accounted for the activity of the neurons, we computed cross-validated negative log-likelihood ratios by normalizing the negative log-likelihoods of single variable models to that of the null-model (Methods). For each of the behaviors considered, and in both pellet reaching and open field tasks, we found that neural responses in PPC and M2 were better predicted by performed behaviors compared to a model with only the constant term (i.e. the mean firing rate; Fig. 5 *A* and *B*). We also noted that the proportions of neurons that were stably tuned to task-dependent behaviors such as grasping (10% in PPC and 24% in M2) and eating (7% in PPC and 6% in M2) fared better than those with task-independent behaviors, such as grooming or rearing. Predictions based on observed behaviors, on the other hand, were in all cases worse than the null-model (Fig. 5 *A* and *B*), which was contrasted strongly by the significant improvement in model performance for the observers’ own movements.

**Fig 5.**
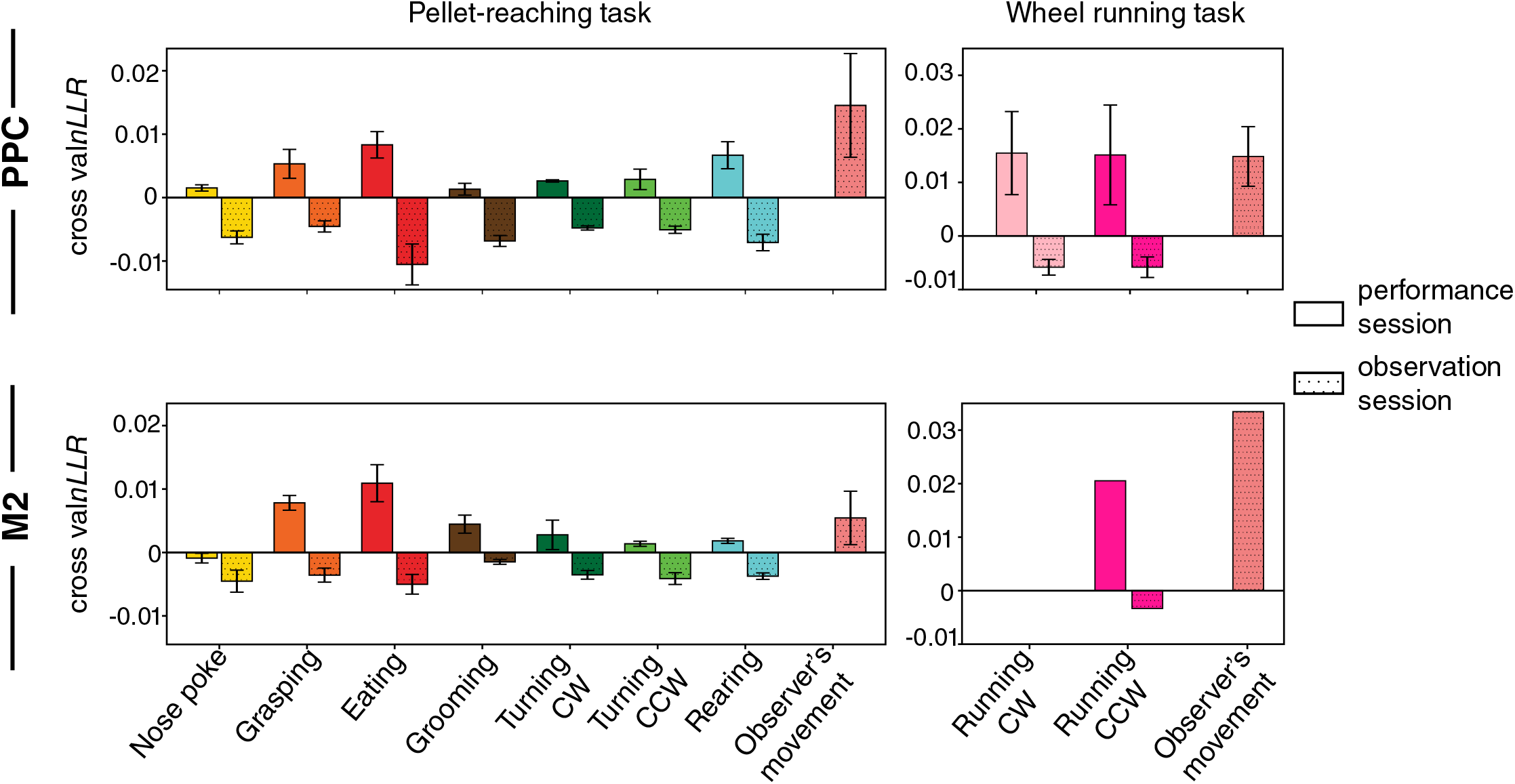
Bar plots show the cross-validated negative log-likelihood ratios for single behavior Bernoulli generalized linear model of calcium events from neural populations in the PPC (*Top panels*) and M2 (*Bottom panels*) of mice during pellet-reaching task (*Left panels*) and wheel running task (*Right panels*). Hand-labeled behaviors from performance sessions are shown as empty boxes while behaviors from observation sessions as hatched boxes. Bars represent the mean ± SEM over animal subjects (pellet-reaching task: 4 mice for PPC and 4 animals for M2; wheel-running task: 3 mice for PPC and 1 mouse for M2).

## Discussion

The results of our study demonstrate that PPC and M2 were reliably modulated by the execution of various natural behaviors in both pellet-reaching and wheel-running tasks, which was juxtaposed sharply by the low number of neurons responding to observed behaviors, which neither exceeded chance levels nor aided in predicting neural activity. Our analysis was inspired by exploration of dimensionally reduced network state dynamics across task conditions, which revealed that population activity in both brain areas was more structured during performance than observation of behaviors. We note that the behavioral clusters in the dimensionality-reduced manifold of performance sessions were not fully separated, which could suggest that the population vectors do not lie completely on a two-dimensional nonlinear manifold, that other tunable parameters of UMAP were not ideally chosen, or that variables which we did not measure, such as posture or decision-making, bind separate behaviors more closely together. In contrast, action observation did not elicit any appreciable structure in population activity. This led us to perform a GLM analysis which confirmed that action observation does not predict neural activity. In fact, the bodily movement of the observers was the most influential factor in the statistical model, which was consistent with results from the performance sessions, and could have been part of a larger wave of neural activation in the brain, as described in recent work in head-fixed animals (36,37).

The fact that the animals were freely moving when performing the tasks allowed us to measure how cells responded to a variety of actions, revealing new features of behavioral coding in both PPC and M2. First, just under 15% of cells in both areas stably represented more than one behavior (Fig. S2), and cells coding for different behaviors were intermingled anatomically. This indicates that cell ensembles in PPC or M2 are apt to participate in more than one behavioral representation, though any overarching organization of tuning based on somatotopy (30), posture (38) or ethological organization (31, 39) was not apparent at the microscales at which we were imaging. Furthermore, while the exact proportion of represented behaviors varied per animal, we generally found turning represented strongly in both PPC and M2, while rearing was more prevalent in PPC and grasping was stronger in M2. In both areas, however, eating was the best predictor of population calcium events in the GLM (Fig. 5), despite that it was coded by comparatively few neurons. Since this predictability could not be attributed to the over-expression of eating epochs relative to other behaviors (Fig S2), it could reflect the salience of the consumptive behavior. It could also imply a population coding strategy where increased single-neuron selectivity compensates for the small population size or, conversely, that a small population size is all that is used because the neurons are strongly tuned (40). On the whole, the heterogeneous response selectivity of cells across distinct behavioral categories is consistent with previous work on multisensory coding and decision making in the rodent PPC (41, 42), while the absence of spatial clustering for similarly tuned neurons is consistent with the dispersed anatomical organization of orientation tuning in primary visual cortex (43), and olfactory coding in the piriform cortex (44).

As for mirror neurons, they have been best characterized across primate species in pre-motor cortex and PPC which, together, comprise the parieto-frontal network (2, 4, 45, 46). This network supports several functions required for goal-directed behavior including sensorimotor transformations, action planning and decision making (47, 48). Although it was long thought that rodent brains lacked the prerequisite complexity to subserve higher cognitive functions, a growing body of work shows that both rats and mice exhibit accomplished performance in sensory-motor tasks such as virtual navigation (49) and evidence-based decision making (20, 50), and they show stimulus history effects (51–53). In terms of anatomy, although PPC and M2 are considerably less elaborate in mice than primates, there are several features common to both species which could support action recognition, including strong input from higher visual areas (54, 55) and dense reciprocal connectivity between PPC and M2 (56–59). Given the anatomical and functional similarities, we reasoned that neurons in the rodent PPC-M2 circuit might exhibit mirror-like responses to the observation and execution of the same actions, and were surprised by the effective absence of observational tuning in both areas.

To our understanding, there are at least two possibilities why this could be the case. One is that the range of behaviors considered was of the wrong kind to elicit mirror responses in rodents. While the pellet reaching task encapsulated the grasping and eating behaviors which evoked mirror neuron activation in primates, it also allowed for the expression of several other natural behaviors, such as grooming and rearing, and wheel running was strongly encoded, particularly in PPC. The absence of observational responses in such tasks suggests that mice may be a species where representations of observed and performed actions do not converge on the same neurons, at least not in the PPC-M2 network, and their capacity for sensorimotor observational learning (13, 14) may depend on non-mirror associative mechanisms. This contrasts with affective learning paradigms, where, for example, mirror-like responses have been shown for pain in the anterior cingulate of rats (19). Thus, distinct anatomical pathways might utilize different neural mechanisms to support different forms of social learning. Another possibility for our findings is that observed actions are encoded in areas outside or upstream of where we imaged. For example, extrastriate areas AL and RL receive the same, if not more, input from V1 as the more medial regions we imaged in PPC, and they project to frontal motor cortices, and could be potential targets for similar experiments in the future.

If the cortical motor system in mice indeed lacks mirror neurons it could also have implications for the evolutionary lineage of sensorimotor mirroring (60). To date, such a phenomenon has been shown in songbirds (6), new (46) and old world monkeys (2, 45), and humans (5, 61). This variety of species raises questions about the phylogenetic development of the capacity for mirroring, and the systems supporting these functions. For example, neurons jointly encoding the vocalization of self and others were found in the telencephalic nucleus HVC of swamp sparrows and zebra finches (6, 62), while audio-vocal mirror neurons were shown in the human inferior frontal gyrus (63). Though avian and primate circuits are not structurally homologous, the question of whether the capacity for mirroring evolved independently or originated in a common ancestor has remained open. Our results suggest the former scenario. Unlike sensorimotor mirroring, vicarious responses for affective states, such as disgust or pain, have been reported in corresponding areas in humans (64, 65) and rodents (19), which is consistent with conserved mirroring involving more ancient sub-cortical systems. This suggests that fundamentally different neural computations may support emotional vs. sensorimotor learning, at least in rodents, and that the likelihood of finding mirror neurons within a system will vary depending on the species in question.

## Materials & Methods

### Subjects and virus injection

All procedures were approved by and in accordance with the Norwegian Animas Act and the European Convention for the Protection of Vertebrate Animals for Experimental and Other Scientific Purposes. Experimental mice were 3 to 7 month old wild type C56BL/6 females (6 from Taconic Bioscience, 2 from The Jackson Laboratory), individually housed on a 12 hr inverted light/dark cycle with ad libitum access to food and water. Surgeries were performed under sterilized conditions and body temperature was maintained at 37°C with a heating pad. Anesthesia was induced using isoflurane mixed with oxygen (5% for induction, 1-1.5% for maintenance) on a stereotactic frame (David Kopf Instruments). Prior to surgery, mice were injected with analgesics subcutaneously (Metacam 1 mg/kg, Temgesic 0.1mg/kg weight) and with a local anesthetic (Marcain 0.5mg/ml) under the skin surface above the skull before making an incision. Following the initial induction and drug administration, the dorsal surface of the head was shaved and ophthalmic ointment was applied to the eyes. The incision area was scrubbed with cotton swabs dipped in 70% Ethanol followed by betadine (2 x each), and a small incision was made along the midline. All measurements were made relative to bregma for virus and prism probe implant surgeries. A craniotomy (1.2 x 1.2mm) was made and each animal was injected with 300nl of AAV1.Syn.GCaMP6m.WPRE.SV40 (University of Pennsylvania Vector Core; item # AV-1-PV2823) at multiple locations in the right hemisphere of the posterior parietal cortex (AP: −1.95, ML: 1.5, DV: 0.35 and 0.7; AP: −1.95, ML: 1.9, DV: 0.35 and 0.7mm relative to bregma) or secondary motor cortex (AP:+0.5, ML: 0.5, DV: 0.5; AP: +0.2, ML: 0.5, DV: 0.5mm relative to bregma) using a Nanoject II Injector (WPI, USA), delivering virus at a rate of 35nl per min with a controller (Micro4; WPI). The glass injection pipette was left in place for 10 min postinjection, after which it was slowly withdrawn. Following the viral injections, the craniotomy was filled with Kwik-Sil silicone elastomer (WPI) and the incision was closed with nylon sutures. After surgery, mice were kept in a heated chamber until they regained consciousness and began moving.

### Prism probe implantation

One week post virus-injection, a 1mm diameter gradient refractive index lens (GRIN) attached to a prism (Inscopix) was lowered stereotaxically into the craniotomy at a rate of 10μm/s while the tissue was treated constantly with saline to minimize desiccation. The prism lens was positioned 1.2-1.3mm deep and 0.15 −0.2mm away from the injection site. Lens implants were secured to the skull with a thin layer of Kwik-Sil silicone elastomer, followed by a thick layer of adhesive cement (super-bond C&B, Sun Medical). The lens cuff was filled with Kwik-Cast (WPI) for protection during a 1-2 week interval to allow for viral expression. A custom made head bar was cemented to the skull with dental acrylic for head fixation in behavioral experiments.

Once viral expression was confirmed, mice underwent anesthesia to secure a baseplate (Inscopix), which was cemented on the prism probe to support the connection of the miniaturized microscope during in vivo imaging under freely moving conditions. During the procedure, a baseplate was attached to the miniature epifluorescence microscope (nVista HD, Inscopix) and stereotaxically positioned to a desired focal plane with the help of visible landmarks (GCaMP6m-expressing neurons and blood vessels) using 20-30% LED power, a frame rate of 5Hz and digital gain of 4. Once the focal plane was identified, the microscope and baseplate were raised by ~50μm to compensate for shrinkage of the adhesive cement, and were subsequently fixed in place using the same compound, followed by a thin layer of dental acrylic mixed with black carbon spherical powder (Sigma Aldrich) to minimize the light interference of the imaging field. The baseplate was covered with a protective cap (Inscopix), and imaging began within 1-2 days.

### Behavioral training and recording

Animal training. Pairs of sibling animals were used in all experiments, and were housed together for one week prior to the start of training. During this period, each animal was habituated to the experimenter and handled extensively on a daily basis. Subsequently, mice were housed individually and food restricted to maintain 90% initial body weight throughout the training period. They were trained daily for 7-10 days in a modified version of the pellet reaching task (29). The chamber used for the task was built from clear plexiglass (3mm thick, 20 x 8.5 x 15cm) with a rectangular cylinder attached externally through which food pellets were delivered (Fig.1). After one day of habituation to the box with no pellets, animals underwent 2 stages of task acquisition; shaping and training. During shaping (2 days, 2 sessions per day), mice were presented with multiple chocolate pellets (20mg per pellet, TestDiet) in the reaching compartment to reinforce reaching behavior. During the subsequent training period (5 days, twice per day), a single pellet was placed in the reaching compartment and the animals’ performance was monitored during 15 min sessions. In this task, each mouse learned spontaneously to turn in a circle in place to elicit pellet delivery (leading to a turn-grasp-eat motif), though this was not explicitly shaped by reinforcement. Trials in which animals retrieved the food pellet with their tongue were excluded from the analysis. Experiments began once mice exceeded 40 successful trials in at least 2 consecutive sessions.

Following head bar placement, the same cohort of animals was gradually habituated to head-fixation over an 8-10 day period. First, they were allowed to move freely in and out of a 4.5 cm diameter acrylic tube, and were subsequently head-fixed with their body in the tube for 15 min. Over 7 days this was increased to 45 min until body movement was minimal. Finally, animals were habituated to head fixation while another conspecific performed the pellet reaching task in front of them. This process typically required ~10 days.

After the pellet reaching task, mice were placed in a wall-less, open, squared arena (30 x 30 cm) with a running wheel, and allowed to behave freely during 20 min sessions. The animals were pre-trained until they exhibited full coverage of the arena, and the same cohort was head-fixed in the tube and, alternately, performed or observed siblings perform the open field task.

Behavioral recording setup. The animals’ behavior was recorded with 5 high-resolution, near-infrared (NIR) cameras (4MP, 100fps, 850nm; Simi Reality Motion Systems GmbH, Germany): one capturing both the performer and observer, one solely on the observer and three exclusively on the performer. The cameras were angled to minimize redundancy of view, and infrared illumination was aided by 8-10 additional NIR LED lamps (850nm, 48 LEDs each; Banggood). All experiments were performed in dim visible light with the experimenter hidden from the view of the animal.

### Pupil measurements

To control for changes in arousal state and neural responsiveness during observation sessions (Reimer et al., 2014), variations in pupil size were measured for 3 mice using close-up video from the camera positioned specifically on the observer, with additional NIR (850nm) illumination of the left eye (Fig. S6). ImageJ software (NIH, version 1.52e) was used to trace a region of interest (ROI) at the lateral edge where the pupil, which was black, met the lighter-colored sclera, which changed dynamically when the pupil dilated or contracted (as in (66)). The mean pixel intensity of the ROI was registered as a negative number that was closest to zero (i.e. largest) when the pupil was dilated maximally, and was most negative when the pupil was contracted (Fig. S6). For each mouse, a binary threshold was determined that captured periods when the pupil was contracting to the smallest size; this was used to flank epochs when the pupil was most contracted, typically when animals were quiescent and motionless.

### Behavioral labeling

Videos were decompressed and downsampled by a factor of 5 (except for one animal which had a 25fps image acquisition rate) to reduce file size and match calcium imaging sampling frequency. The videos of several behavioral sessions were reviewed closely to determine which behaviors were sufficiently frequent and reliable to label manually, including task-specific (e.g. grasping a pellet) and non-specific (e.g. rearing) behaviors. The behaviors were manually labelled using a Jython-based, custom-developed graphical user interface (GUI). For each recording session, videos with different fields of view (with at least one of the performer and one of the observer) were loaded into the GUI, and two experimenters scored behaviors from the same sessions frame by frame. The behaviors used for subsequent neural analyses included nose poke, grasping, eating, grooming, turning (with clockwise and counter-clockwise turning separated) and rearing (Movie S2). In the open field we only quantified wheel-running behavior, but again discretized clockwise and counterclockwise directions. We also labeled epochs when observer animals moved their limbs or bodies during the observation experiments, allowing us to measure neuronal activity during observer movement.

### Calcium imaging

One photon imaging of intracellular calcium activity was acquired at a rate of 20-25Hz, with LED power set to 20-30% and a gain of 1; the same image acquisition parameters were maintained for a given set of sessions (4 x 10 min) to allow for comparison of neural activity (27). Calcium imaging timestamps were synchronized with the behavioral recording system for offline behavioral analyses. Synchronization was done using the nVista DAQ box (Inscopix), which enabled triggering of external hardware (behavioral recording system; Simi) using a TTL system. GCaMP6m-expressing C57BL/6 mice were imaged while performing the pellet reaching task (2 x 10 min), and again while observing the task (2 x 10 min) while head-fixed. The following day, the same animals were imaged while freely exploring the open field with the running wheel (2 x 10 min), and again while head-fixed, observing a conspecific doing the same (2 x 10 min).

### Image processing

Fluorescence movies were processed using Mosaic Software (v.1.1.2, Inscopix). Raw videos were spatially downsampled by a factor of 4 to reduce file size and processing time; temporal downsampling was not applied. Dropped frames were isolated and interpolated, and the movies were cropped to remove regions lacking cells. For pellet-reaching and open field experiments, performance and observation recordings of the same task were concatenated to generate a single 40 min recording. Motion artifacts were corrected using a single reference image (typically obtained by drawing a border around a large blood vessel or selecting bright neurons) using the Turboreg image registration algorithm within Mosaic software. The movies were further cropped to remove post-registration black borders.

### Fluorescence trace extraction

Motion-corrected, cropped recordings were saved as .tiff files for subsequent signal extraction using the constrained non-negative matrix factorization algorithm for endoscopic recordings (CNMF-E) (67). CNMF-E was designed to isolate large fluctuations in background fluorescence and facilitate the accurate extraction of cellular signals by simultaneously denoising, deconvolving and demixing one photon calcium imaging data. The CNMF-E framework can be summarized by the following steps: (1) initialize the spatial and temporal components of all neurons without explicit estimation of the background, (2) approximate the background given the activity of all neurons, (3) update spatial and temporal components by subtracting background from the raw image using alternating matrix factorization, (4) delete neurons and merge neurons with high temporal correlations, (5) repeat steps 2-4 (for quantitative detail see Zhou, et al., 2018). Similar parameters (gSig = 3, gSiz = 13, mincorr = 0.9) were used across different data sets to extract fluorescence signals. After calcium signal extraction with CNMF-E, fluorescence traces were deconvolved to approximate relative firing rates in each imaging frame using ‘Online Active Set methods for Spike Inference’ (OASIS) (68). For this, the fluorescence data was modelled using an autoregressive (AR(1)) process due to the fast rising time of calcium. The decay time of the calcium signal (g hyperparameter) was estimated from the autocorrelation, and the optimized g hyperparameter was set to 0. Lastly, a strict threshold of 5 standard deviations from the mean event was used for further calcium event estimation. All subsequent analyses used the inferred calcium events to minimize the effect of decay kinetics of calcium signals.

### Signal-to-noise ratio

A signal-to-noise ratio (SNR) analysis was performed to estimate the quality of the deconvolved output relative to raw traces. Every raw trace value in the interval spanning one second before to seven seconds after a registered calcium event (to accommodate the sharp rise and slow decay of the calcium signal) was considered as signal, and everything outside that range was considered as noise. The SNR was defined as the ratio of the mean of the traces related to calcium events and the standard deviation of the noise. Any cell that failed to exceed or match the SNR minimum value of 3.5 for all sessions was discarded from further analyses.

### Behavioral tuning and shuffling

Calcium event rates were calculated for each cell during each behavior by dividing the total number of events within a behavior by the total time spent in that behavior (in seconds). The calcium event trains were then offset by a random interval between 20 and 60 sec one thousand times, and event rates for each behavior were recalculated for each permutation, generating a shuffled distribution. The observed firing rates were z-scored relative to the shuffled distribution, and a cell was considered significantly tuned if its z-scored rate was 2 standard deviations above its shuffled mean during a given behavior. Only cells meeting this criterion for two of the same type of session were considered stably tuned. During observation sessions, the observers’ body movements were registered in addition to the behavior of the performer. Cells tuned to the observer’s movement in any session were discarded from the analysis as potentially showing tuning to observed actions.

### Peri-event time histograms

Calcium events were binned in 200 ms windows relative to the onset of a given behavior, converted into rates and convolved with a Gaussian kernel with a width of 1 bin. Behavioral epochs shorter than 100 ms were excluded from the analysis. For each bin, the mean and the standard error of mean were calculated over epochs. After averaging over epochs, each cell was normalized to its peak rate and cells were ordered according to the magnitude of their z-scored rate in the first performing session (P1).

### False positive estimations

For either brain area, we assessed whether the number of stably tuned neurons across different conditions (performance, observation and matched) was statistically different from chance (i.e. false positive) rates. The false positive rate was estimated empirically by swapping behavioral labels between two sessions of the same kind (e.g. O1 and O2) and re-computing calcium rates for each behavior, thus determining the “false” proportion of stably tuned cells across all animals. The significance of the difference between the distributions of true and false positive proportions was determined with the Mann-Whitney U test.

### Correlation matrices

To assess the predictability of representations across different session types, data from all animals within a region were pooled and significantly tuned cells for each behavior (e.g. rearing) in each session (e.g. P1) were selected. A given z-scored calcium rate series (e.g. all rearing cells in P1) was then correlated with the series of z-scored rates of all the behaviors in all the other sessions (e.g. all grooming cells in O1).

### Cell registration

To identify discrete states in neuronal population activity using dimensionality reduction (Uniform Manifold Approximation and Projection, UMAP; Mclnnes et al., 2018), the stability and identity of cells across all sessions were first confirmed using methods recently published by Sheintuch et al. (69), which uses a probabilistic approach to register the spatial location of cells across sessions. After extraction of spatial components of the imaged data for each recording, spatial footprints were loaded into a graphical user interface (GUI) provided by Sheintuch et al. (2017) for further alignment and characterization of the similarity measure. For this analysis, a pixel value of 2.3μm, maximal distance of 15μm (due to sparsity) and Psame threshold of 0.95 (to be conservative) were used.

### Dimensionality reduction

The fast, non-linear dimensionality reduction algorithm, UMAP, was applied to visualize the high-dimensional neural state space using a lower-dimensional manifold while preserving high-dimensional local and global structures. To do this, cells were first registered across a total of 60 minutes of combined pellet reaching and open field recordings (described in “Cell registration”) to ensure similarity. The calcium event trains were then binned to the resolution of the imaging sampling rate (20Hz for PPC, 25Hz for M2) and the activities were convolved using a Gaussian kernel with a width of 2 bins. Neural data was downsampled to every 2 bins, then further downsampled by keeping only the time points when >10% of the population for performing sessions and >5% for observing sessions had non-zero convolved events. Dimensionality reduction with UMAP was performed assuming a Manhattan distance metric, and the parameters (n_neighbors=5, min_distance=0.5, spread=1.0) were kept the same for all neural data sets.

### Dunn Index

The compactness of the behavioral clusters (i.e., cluster of time points corresponding to the same hand-labeled behavior) in the dimensionality-reduced representation was assessed using the Dunn Index (DI) (70). To this end, the centroids were first calculated for every behavioral cluster. Distances between each point within a behavioral cluster and the cluster’s centroid (intra-cluster distances) and the distances between centroids of different clusters (inter-cluster distances) were measured. The DI was then calculated as the ratio between the minimum inter-cluster distance and the maximum intra-cluster distance (as defined above). The DI provides a measure of overall clustering quality, i.e., a high Dunn index corresponds to tight clustering in the data.

### Generalized linear model

For performed behaviors, the neural calcium event data from performing sessions was fitted with generalized linear models (GLMs) to determine whether a given performed behavior explained the calcium events better than the neurons’ mean calcium events rate. To do this, the events were binned to the resolution of the imaging sampling rate (20Hz). The calcium event data were then fitted with a Bernoulli GLM (35) assuming the neurons were independent. Each GLM contained a parameter corresponding to a hand-labeled behavior (nose poke, pellet grasping, eating, grooming, turning CW, turning CCW, rearing, running CW or running CCW) as well as a constant term. The likelihood of the data given each of the models was maximized across 10 folds of the data. Calcium events recorded from each neuron were also fitted with a Bernoulli GLM (which we call the null model) with only the constant term, which corresponded to the neuron’s mean calcium event rate. The out-of-sample likelihood was calculated for each fitted GLM. The cross-validated negative log-likelihood ratio (cross-val nLLR) was then calculated as the difference between the out-of-sample model likelihood, which was obtained from the GLM with a parameter attached to a hand-labeled behavior, and the out-of-sample null model likelihood, which was from the GLM with only the constant term, normalized over the out-of-sample null model likelihood and averaged over 10 folds of the data.

For observed behaviors, the neural calcium event data from observing sessions were also fitted with Bernoulli GLMs to determine whether a given observed behavior could account for the calcium events beyond what can be explained with the observer’s own behavior (i.e., body movement). The cross-val nLLRs were calculated as with performed behaviors, but with the out-of-sample model likelihood obtained from the GLM with parameters attached to an observed behavior and to body movement, plus the out-of-sample null model likelihood from the GLM with a parameter attached to only the body movement.

### Spatial clustering of behaviorally selective neurons

To calculate the spatial distribution of significantly tuned neurons, each neuron’s centroid location was first identified using TrakEM2 software (71). To do this, neurons identified as responsive to any given behavior were stacked together in ImageJ, and image stacks for each behavior were averaged to obtain a single image with the physical locations of tuned neurons. These images were subsequently loaded into TrakEM2 and the position of each cell in each image was manually traced as a circle. The XY location of each circle was calculated to obtain the position of each cell in each animal. Next, Euclidean distances between stably tuned cells in the imaging field for each mouse were calculated. To evaluate spatial clustering of cells based on their behavioral correlates, the Dunn Index (DI; see Dimensionality reduction: Dunn index) of each animal’s recorded dataset was compared against the distribution of DIs generated from shuffled data. A behavioral cluster was defined as the cluster of cells that was stably tuned to a given behavior. The shuffled distribution of DIs was obtained by randomly permuting cell IDs one thousand times and recalculating the DI for each permutation.

### Anatomical verification of imaging locations

For perfusions, animals were anaesthetized deeply using isoflurane (5%) and subsequently injected with sodium pentobarbital (200 mg/kg; intraperitoneal injection) and transcardially perfused using ~25 ml saline followed by ~50 ml of 4% paraformaldehyde (PFA). Each mouse was decapitated and the brain was removed carefully from the skull. Brains were kept in 4% PFA at 4 °C overnight, then transferred to 2% dimethyl sulfoxide (DMSO: VWR, Radnor, PA) solution for cryoprotection for 1-2 days. The brains were cut in coronal sections in 3 series of 40μm on a freezing sliding microtome (HM-430 Thermo Scientific, Waltham, MA). The first series was mounted directly onto the superfrost slides (Thermo Scientific) to perform Nissl-staining for delineation purposes. The remaining series of sections were collected in vials containing 2% DMSO and 20% glycerol in phosphate buffer (PB) and stored at −20° C until further usage.

For immunohistochemical staining, the second series of sections was used to visualize GCaMP6m viral expression. The brain sections were first rinsed 3 x 5 min in PBS on a shaker, incubated in blocking buffer (PBS plus 0.3% Triton, 2 x 10 min), followed by incubation in primary antibody solution (rabbit anti-GFP, 1:1000, ThermoFisher Scientific, A-11122, in PBS and 0.3% Triton) overnight at 4 °C. Sections were further washed in PBS containing 0.3% Triton and 3% bovine serum albumin (BSA; Sigma Aldrich) for 2 x 5 min at room temperature (RT), and subsequently incubated in secondary antibody solution (AlexaFluor 488-tagged goat anti-rabbit Ab, 1:1000, ThermoFisher Scientific, A-11008) for 1 h at RT. Sections were washed 2 x 10 min in PBS and mounted on gelatin-coated polysine microscope slides and dried in the dark overnight. Next, sections were treated with Hoechst solution (1:5000; Sigma Aldrich) for 5 min in the dark and immediately rinsed with PBS. Slides were air dried overnight in the dark at RT and cover-slipped using entellan-toluene solution (Merck Chemicals) the following day.

For anatomical delineation of recording locations, all brain sections were digitized using an automated scanner for fluorescence and brightfield images at the appropriate illumination wavelengths (Zeiss Axio Scan.Z1, Jena, Germany). Corresponding Nissl stained sections were used to delineate PPC, M2 and neighboring cortical regions in each animal in accordance with Hovde et al. (2018), the borders of which were copied onto the GFP-stained images in Adobe Illustrator CC 2017. Bregma coordinates were estimated in correspondence with Paxinos & Franklin (72).

## Supporting information

UMAP

Behaviors_pellet_reaching_task

## Acknowledgements

We thank C. Keyser and E. Moser for helpful comments on the manuscript; S. Gonzalo Cogno and H. Obenhaus for fruitful discussions on the analysis. H. Kleven, M. Gianatti, C. Bjørkl i, and H. Waade for technical and I.T. assistance; E. Demirci for assistance with behavioral labeling; M. Witter for helpful discussions on anatomy; S. Eggen for veterinary oversight. This study was supported by the European Research Council (‘RAT MIRROR CELL’, Starting Grant Agreement N° 335328), the Research Council of Norway (FRIPRO Young Research Talents, Grant Agreement N° 239963), the Kavli Foundation, and the Center of Excellence scheme of the Research Council of Norway (Center for Neural Computation).

## Author contributions

J.R.W., T.T. and B.A.D. designed the project; T.T., B.A.D., K.H. designed aspects of experiments; T.T. and K.H. contributed half the data each. B.A.D., B.M., R.J.C., Y.R. and P.M. designed the analyses, T.T., K.H., R.J.C., and B.M. performed the analyses. T.T., B.M., J.R.W. wrote the paper with assistance from R.J.C., Y.R., K.H., and B.A.D.

The authors declare no conflict of interest.

## Supplementary Figure legends

**Fig. S1.**
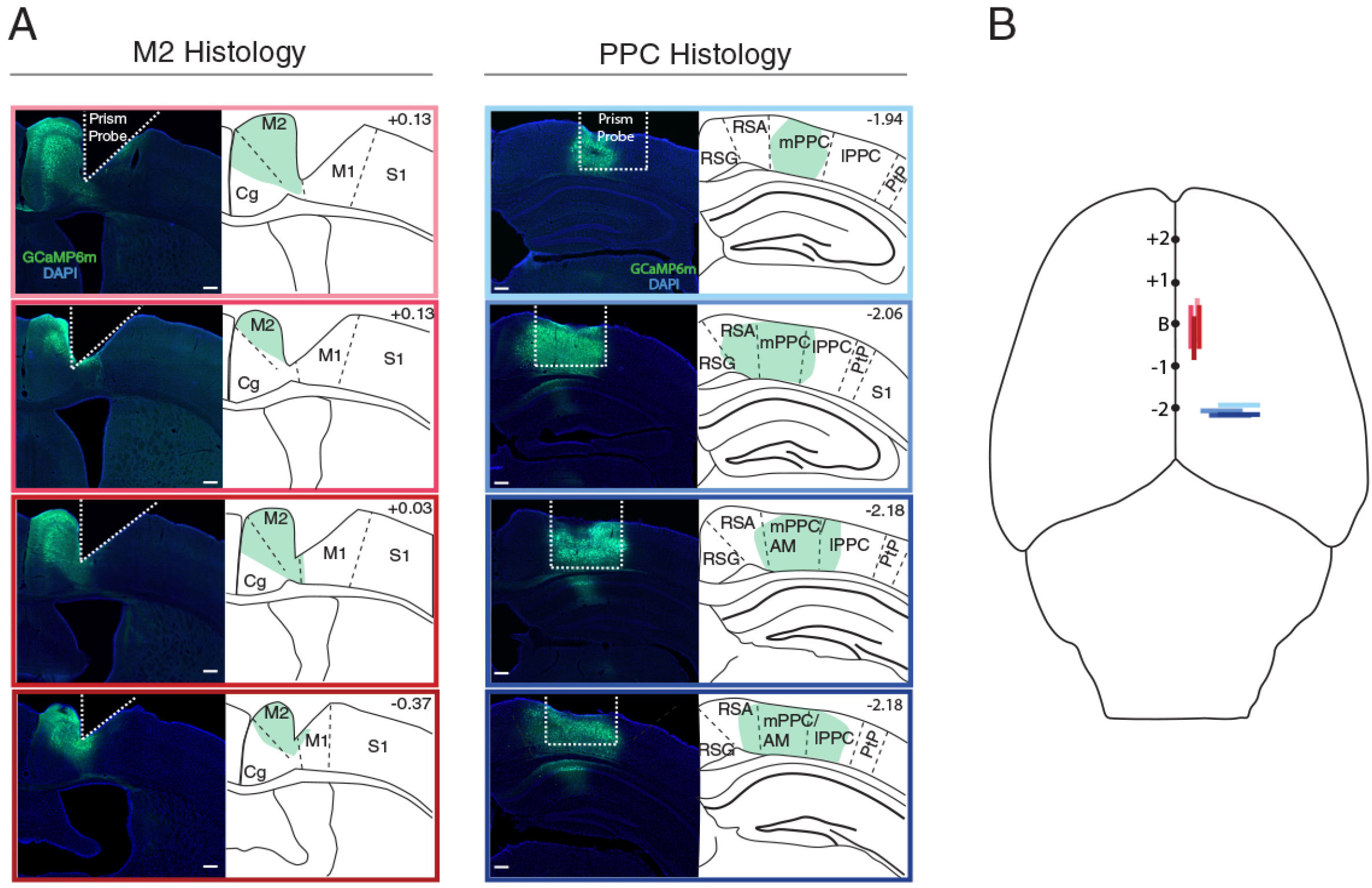
(*A*) (*Left*) Histological sections (40μm) showing GCaMP6m expression in M2, with prism probe locations depicted by the white dashed line. (*Right*) Same, for animals in PPC. In both areas, schematics of the tissue were drawn to show the extent of GCaMP6 expression in green. Anatomical boundaries for PPC, M2 and surrounding regions were established using lamination and cytoarchitectural profiles in adjacent, Nissl-stained sections. Scale bar denotes 200 μm. (*B*) Dorsal view of estimated recording planes in M2 (red rectangles) and PPC (blue) in all 8 animals. Bregma (“B”) is indicated on the midline, and black dots indicate 1 mm.

**Fig. S2.**
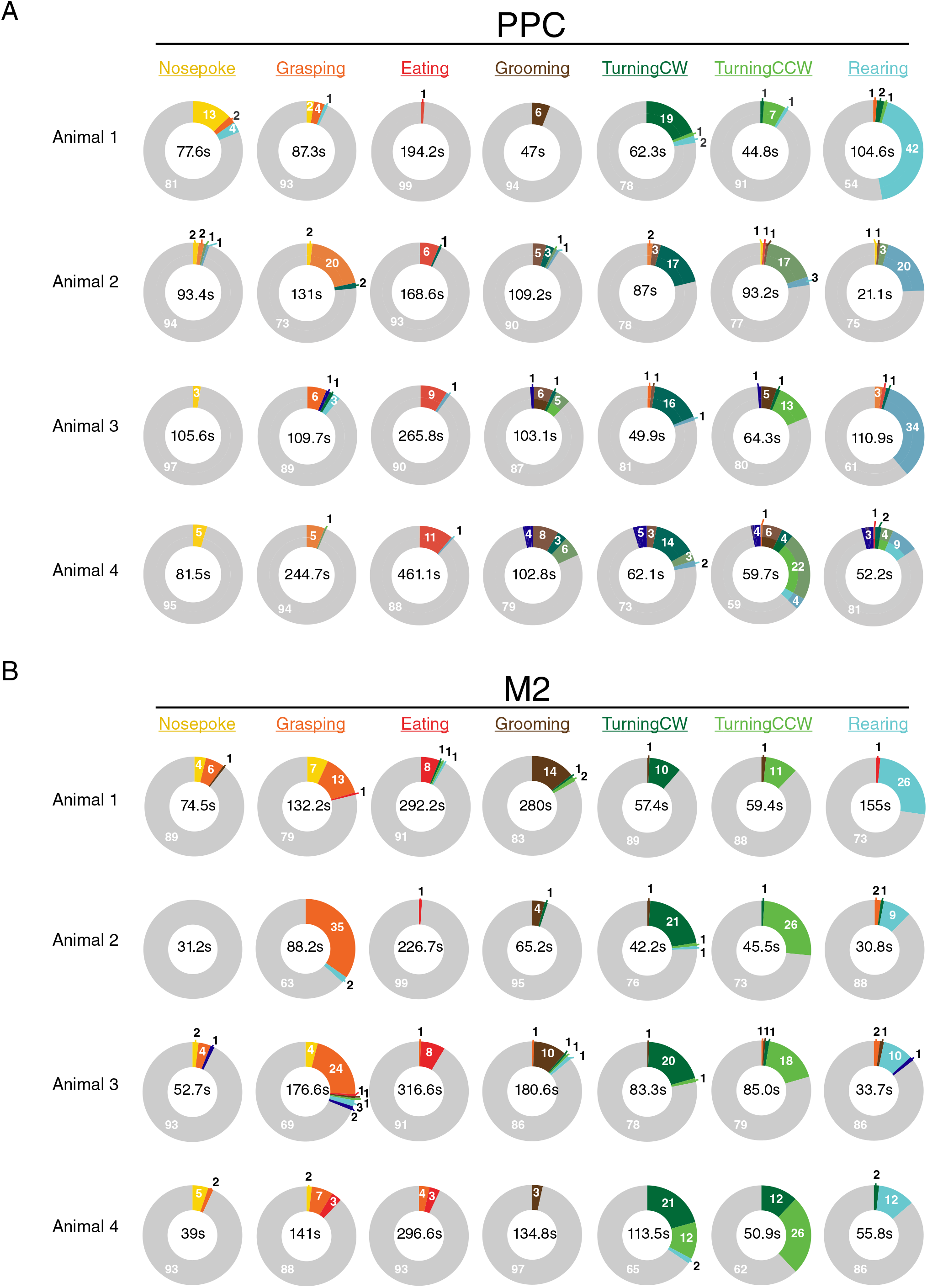
Subsets of cells in PPC and M2 were stably tuned to multiple behaviors. (*A*) Color-coded pie charts show the proportion of PPC neurons significantly tuned to each behavior in the pellet reaching task, with the number of cells in each category written around the ring periphery, and the total time in each behavior (summed across both performance sessions) shown in the center. To display the relative proportions graphically, cells tuned to multiple behaviors (e.g. “Nose poke” and “Grasping”) appear in more than one pie chart. Cells stably tuned to three or more behaviors are denoted by dark blue, while cells not tuned to the behavior of interest are shaded in grey. (*B*) Same as in *A*, but for M2.

**Fig. S3.**
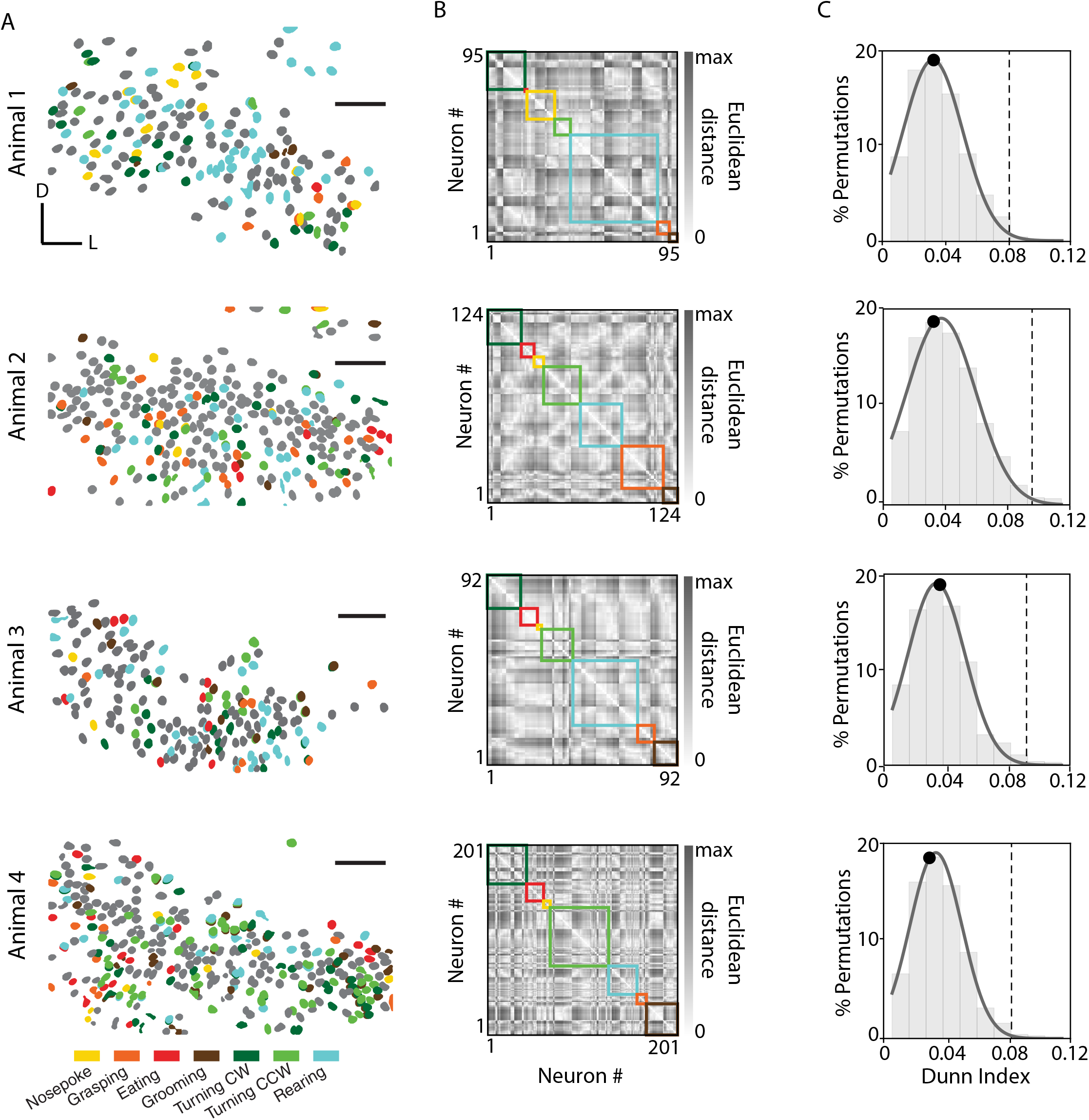
Behaviorally tuned neurons in PPC did not cluster anatomically. (*A*) Cell maps for each animal, color-coded by their behavioral correlates (legend at bottom). Scale bars = 100μm. (*B*) Matrices showing pairwise Euclidean distances between neurons grouped by their tuning preferences (colored boxes); shortest distances are shown in white and longer distances are darker. Functional-anatomical clustering would produce lighter shading within-behavior and darker colors outside. (*C*) The quality of clustering by behavior was quantified using the Dunn index (Methods), which assessed Euclidean distances between cells with similar vs. different behavioral classifications. The distribution of actual intra-vs. inter-cluster distances was compared against a shuffled distribution in which cell identities were permuted, which indicated below-chance levels of clustering in each animal. Dashed lines indicate the 99th percentile of the shuffled distribution; black circles denote the Dunn index value.

**Fig. S4.**
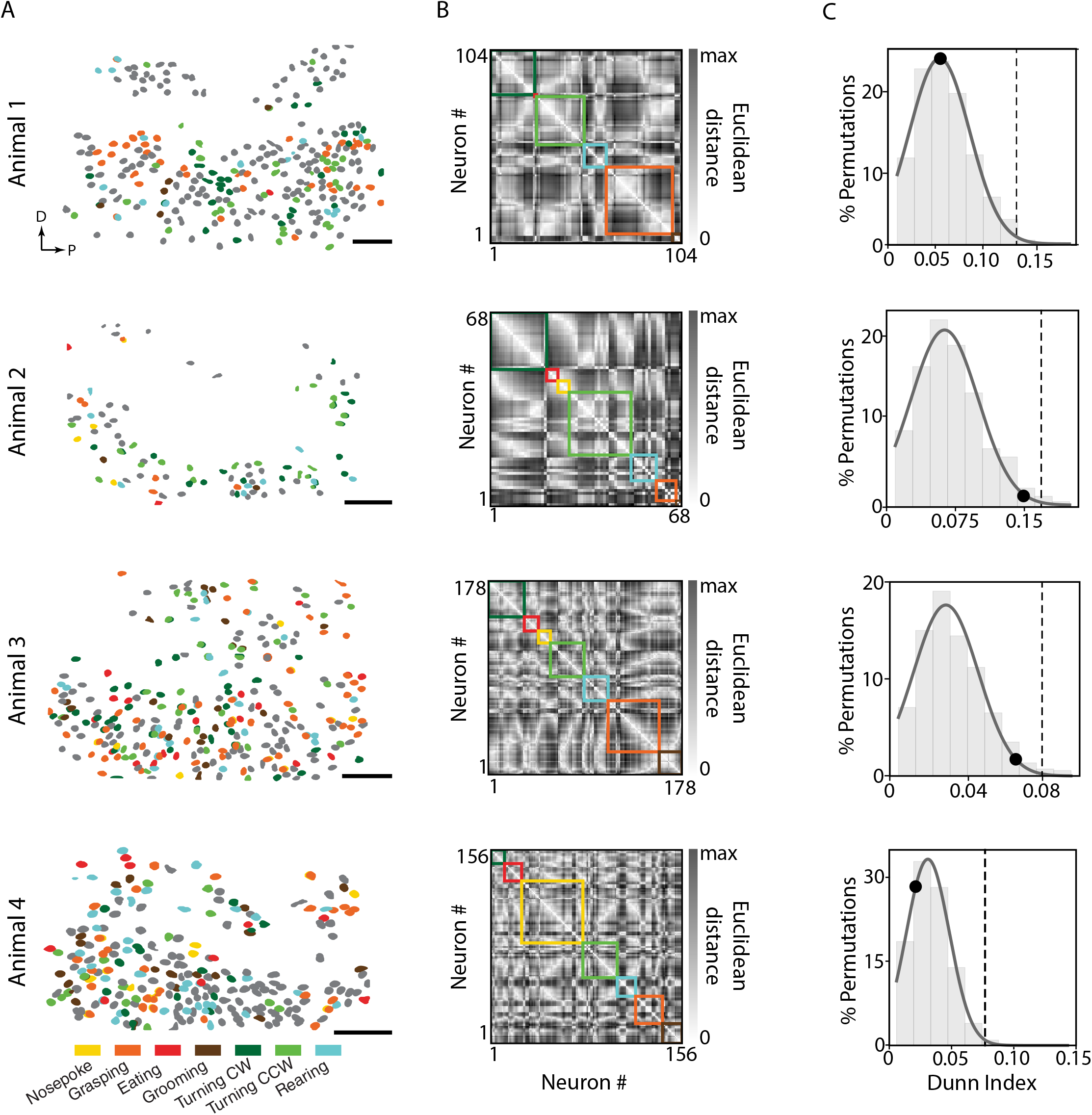
Behaviorally tuned neurons in M2 were not clustered anatomically. (*A*) Cell maps for each animal, color-coded by their behavioral correlates (legend at bottom). Scale bars = 100μm. (*B*) Same matrices as for PPC cells in Figure S3, showing pairwise Euclidean distances between neurons grouped by tuning preferences. (*C*) The quality of clustering by behavior was quantified using the Dunn index (Methods), as with PPC neurons in the previous Supplementary figure; none of the animals showed neural clustering exceeding the 99th percentile of the shuffled distribution (dashed lines); black circles denote the observed Dunn index value.

**Fig. S5.**
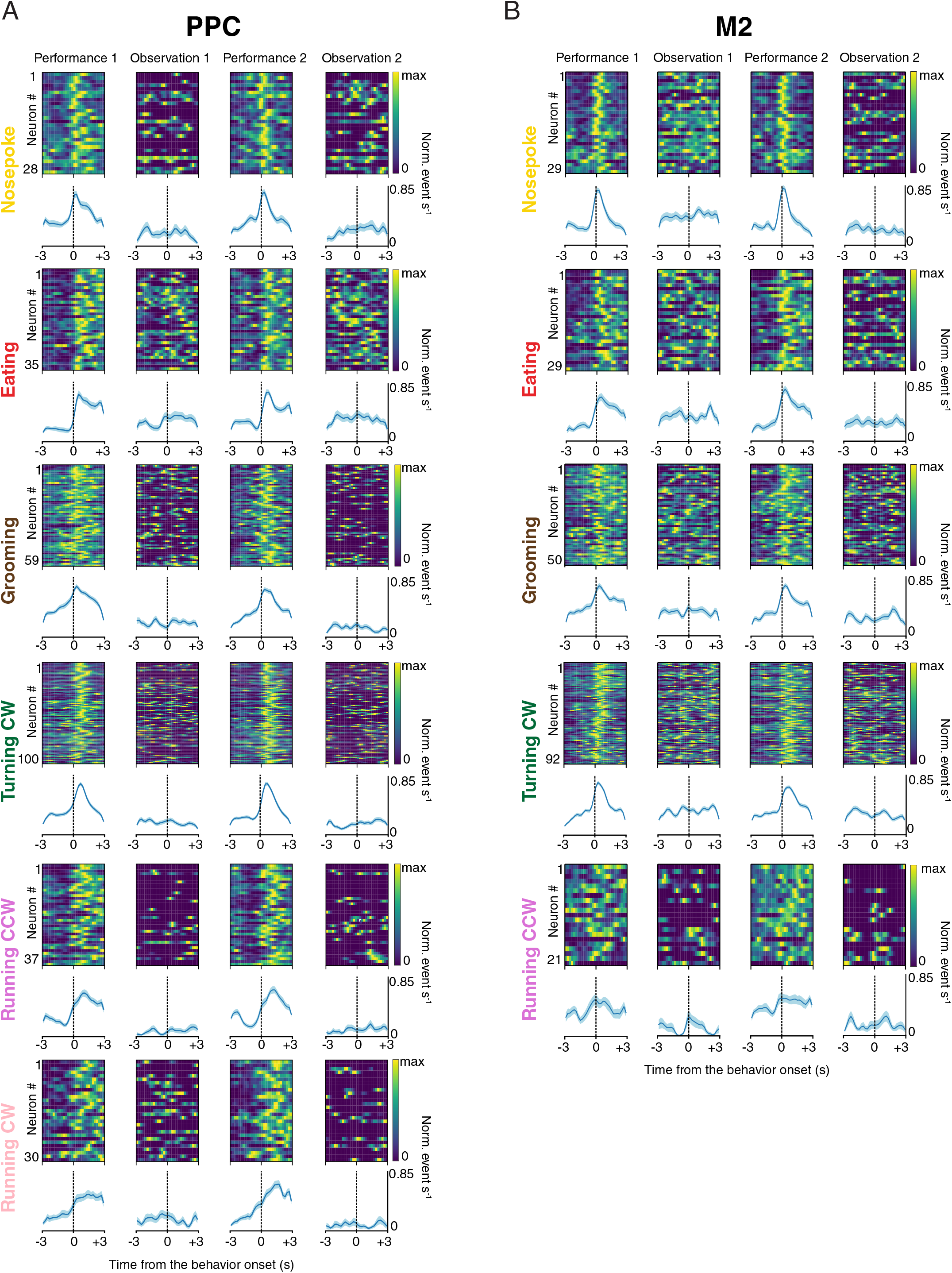
Additional behavioral conditions in relation to Figure 4 comparing PPC and M2 ensemble activation during performance and observation sessions. (*A*) As with Figure 4, PPC cells responded during performed, but not observed actions. (*B*) Same as *A*, but for cells recorded in M2; insufficient data were collected to test for stable tuning for Running CW for recordings in M2, so that condition was omitted. Note that the behaviors here are included in the cross-correlation matrices for performance and observation sessions in Figure 4 *C* and *D*.

**Fig. S6.**
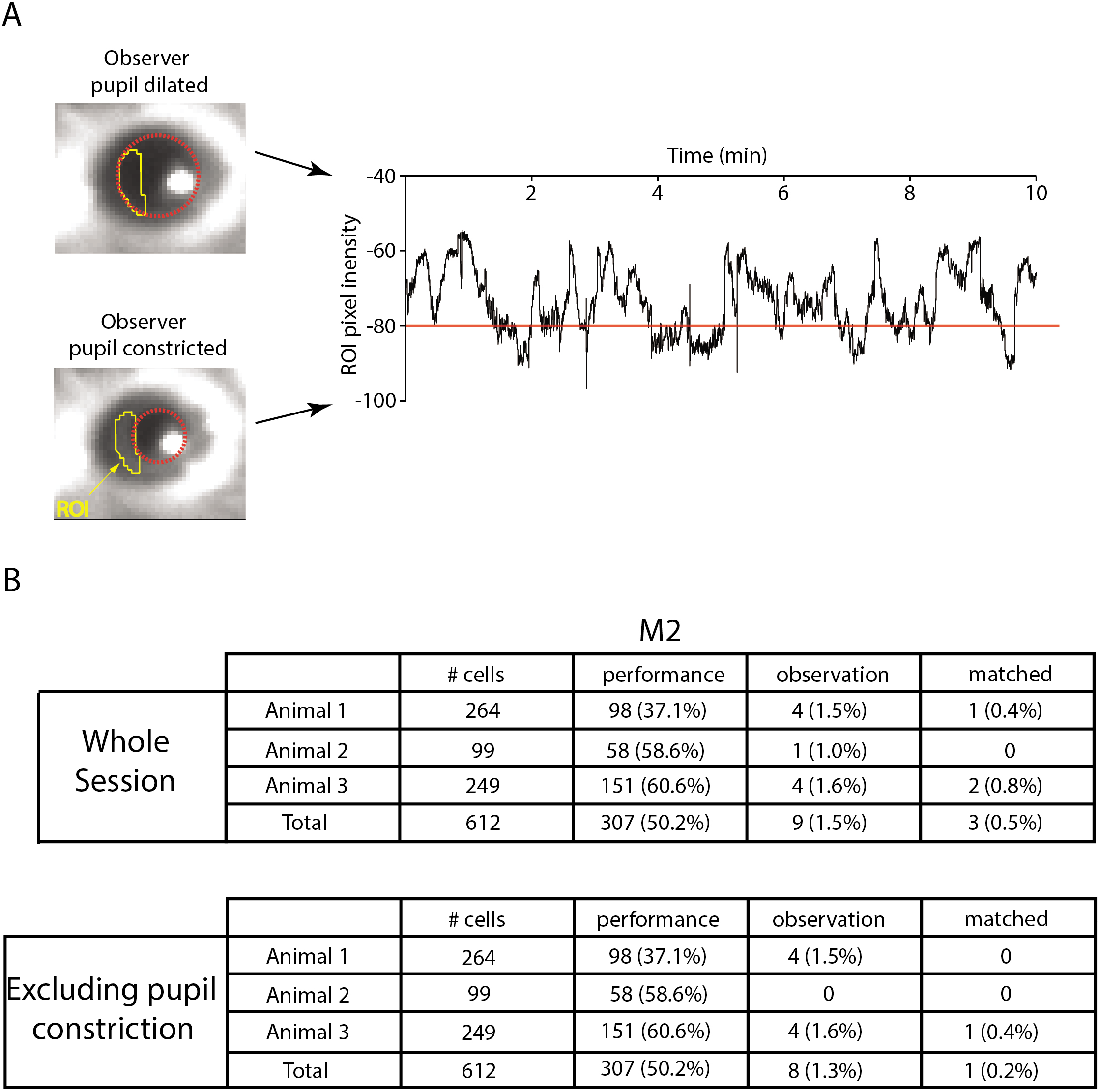
Arousal state did not influence neural responses to observed actions. (*A*) Pupil size was measured as a proxy for arousal state during observation of the pellet-reaching task in three mice with prisms in M2. (*Left*) A region of interest (ROI) was drawn over a close-up video of the eye using ImageJ software, and pupil size (red circle) was reported via pixel intensity inside the ROI (*Right*). For each mouse, a threshold was determined to capture epochs when the pupil was constricting to its smallest size (red line in graph), typically when animals were quiescent and motionless. (*B*) The number of cells with stable correlates for observed behavior was below the false positive rate regardless of whether epochs with small pupil diameter were included in the analysis.

## Supplementary Movies

**Movie S1.** (left panel) The momentary state of neural population activity is indicated by the blue cursor, while the dimensionally-reduced manifold of population activity for the entire session is shown as grey dots. Darker areas correspond to denser regions in the reduced space. Note that the cursor (i.e. the state of population activity) occupies a stable location when the animal performs clockwise and counter-clockwise running, but that it moves unpredictably over the manifold when the same animal observes a cohort running on the wheel. (right panel) Corresponding in-session videos of wheel-running epochs from performance and observation sessions. For display purposes, calcium events were convolved with a Gaussian kernel with width of 5 bins before down sampling and manifold learning using UMAP (Methods).

**Movie S2.** Video showing a side-view of a mouse performing the pellet reaching task. Each behavior included in the neural data analyses is demonstrated in the video.

**Table S1.** Summary of performance and observation tuning in PPC and M2

|  |  | # cells | performance | observation | matched |
| --- | --- | --- | --- | --- | --- |
| Pellet Reaching | Animal 1 | 177 | 85 (48%) | 1 (0.6%) | 0 |
|  | Animal 2 | 268 | 111 (41.4%) | 4 (1.5%) | 0 |
|  | Animal 3 | 179 | 80 (44.7%) | 3 (1.7%) | 1 (0.6%) |
|  | Animal 4 | 297 | 154 (51.9%) | 7 (2.4%) | 2 (0.7%) |
|  | Total | 921 | 430 (46.7%) | 15 (1.6%) | 3 (0.3%) |
| Wheel-Running | Animal 1 | 149 | 20 (25.5%) | 0 | 0 |
|  | Animal 2 | 337 | 37 (10.9%) | 1 (0.3%) | 0 |
|  | Animal 3 | 238 | 10 (4.5%) | 2 (0.8%) | 0 |
|  | Total | 724 | 67 (9.3%) | 3 (0.4%) | 0 |

M2
|  |  | # cells | performance | observation | matched |
| --- | --- | --- | --- | --- | --- |
| Pellet Reaching | Animal 1 | 264 | 98 (37.1%) | 4 (1.5%) | 1 (0.4%) |
|  | Animal 2 | 99 | 58 (58.6%) | 1 (1.0%) | 0 |
|  | Animal 3 | 249 | 151 (60.6%) | 4 (1.6%) | 2 (0.8%) |
|  | Animal 4 | 240 | 132 (55%) | 4 (1.7%) | 0 |
|  | Total | 852 | 439 (51.5%) | 13 (1.5%) | 3 (0.4%) |
| Wheel-Running | Animal 1 | 216 | 21 (9.7%) | 1 (0.5%) | 0 |
|  | Total | 216 | 21 (9.7%) | 1 (0.5%) | 0 |

## References

1. B. G. Galef, Jr.; Laland, K. N., Social Learning in Animals: Empirical Studies and Theoretical Models. BioScience 55, 489–499 (2005).

2. G. di Pellegrino, L. Fadiga, L. Fogassi, V. Gallese, G. Rizzolatti, Understanding motor events: a neurophysiological study. Experimental brain research 91, 176–180 (1992).

3. V. Gallese, L. Fadiga, L. Fogassi, G. Rizzolatti, Action recognition in the premotor cortex. Brain 119 (Pt 2), 593–609 (1996).

4. L. Fogassi et al., Parietal lobe: from action organization to intention understanding. Science (New York, N.Y 308, 662–667 (2005).

5. R. Mukamel, A. D. Ekstrom, J. Kaplan, M. Iacoboni, I. Fried, Single-neuron responses in humans during execution and observation of actions. Curr Biol 20, 750–756 (2010).

6. J. F. Prather, S. Peters, S. Nowicki, R. Mooney, Precise auditory-vocal mirroring in neurons for learned vocal communication. Nature 451, 305–310 (2008).

7. G. Rizzolatti, L. Craighero, The mirror-neuron system. Annual review of neuroscience 27, 169–192 (2004).

8. M. Iacoboni, Imitation, empathy, and mirror neurons. Annu Rev Psychol 60, 653–670 (2009).

9. J. D. Russo, Observational learning in hooded rats. Psychonomic Science 24, 37–38 (1971).

10. O. Zohar, Terkel, J., Acquisition of pine cone stripping behavior in black rats (rattus rattus). International Journal of Comparative Psychology 5, 1–6 (1991).

11. M. G. Leggio et al., A new paradigm to analyze observational learning in rats. Brain Res Brain Res Protoc 12, 83–90 (2003).

12. M. Kavaliers, E. Choleris, D. D. Colwell, Learning from others to cope with biting flies: social learning of fear-induced conditioned analgesia and active avoidance. Behav Neurosci 115, 661–674 (2001).

13. P. Carlier, M. Jamon, Observational learning in C57BL/6j mice. Behavioural brain research 174, 125–131 (2006).

14. M. T. Jurado-Parras, A. Gruart, J. M. Delgado-Garcia, Observational learning in mice can be prevented by medial prefrontal cortex stimulation and enhanced by nucleus accumbens stimulation. Learn Mem 19, 99–106 (2012).

15. D. J. Langford et al., Social modulation of pain as evidence for empathy in mice. Science 312, 1967–1970 (2006).

16. T. Sakaguchi, S. Iwasaki, M. Okada, K. Okamoto, Y. Ikegaya, Ethanol facilitates socially evoked memory recall in mice by recruiting pain-sensitive anterior cingulate cortical neurons. Nat Commun 9, 3526 (2018).

17. D. Jeon et al., Observational fear learning involves affective pain system and Cav1.2 Ca2+ channels in ACC. Nature neuroscience 13, 482–488 (2010).

18. S. A. Allsop et al., Corticoamygdala Transfer of Socially Derived Information Gates Observational Learning. Cell 173, 1329–1342 e1318 (2018).

19. M. Carrillo et al., Emotional Mirror Neurons in the Rat’s Anterior Cingulate Cortex. Curr Biol 29, 1301–1312 e1306 (2019).

20. T. D. Hanks et al., Distinct relationships of parietal and prefrontal cortices to evidence accumulation. Nature 520, 220–223 (2015).

21. A. M. Licata et al., Posterior Parietal Cortex Guides Visual Decisions in Rats. J Neurosci 37, 4954–4966 (2017).

22. H. Mohan, R. de Haan, H. D. Mansvelder, C. P. J. de Kock, The posterior parietal cortex as integrative hub for whisker sensorimotor information. Neuroscience 368, 240–245 (2018).

23. F. Barthas, A. C. Kwan, Secondary Motor Cortex: Where ‘Sensory’ Meets ‘Motor’ in the Rodent Frontal Cortex. Trends Neurosci 40, 181–193 (2017).

24. J. C. Erlich, M. Bialek, C. D. Brody, A cortical substrate for memory-guided orienting in the rat. Neuron 72, 330–343 (2011).

25. J. R. Whitlock, G. Pfuhl, N. Dagslott, M. B. Moser, E. I. Moser, Functional split between parietal and entorhinal cortices in the rat. Neuron 73, 789–802 (2012).

26. A. Rubin, Sheintuch, L., Brande-Eilat, N., Pinchasof, O., Rechavi, Y., Geva, N., Ziv, Y., Revealing neural correlates of behavior without behavioral measurements. biorXiv https://doi.org/10.1101/540195 (2019).

27. K. K. Ghosh et al., Miniaturized integration of a fluorescence microscope. Nat Methods 8, 871–878 (2011).

28. L. McInnes, Healy, J., Melville, J., UMAP: Uniform Manifold Approximation and Projection for Dimension Reduction. arXiv (version 2 of preprint), arXiv:1802.03426 (2018).

29. T. Xu et al., Rapid formation and selective stabilization of synapses for enduring motor memories. Nature 462, 915–919 (2009).

30. R. D. Hall, Lindholm, E.P., Organization of motor and somatosensory neocortex in albino rat. Brain research 66, 23–38 (1974).

31. D. F. Cooke, C. S. Taylor, T. Moore, M. S. Graziano, Complex movements evoked by microstimulation of the ventral intraparietal area. Proc Natl Acad Sci U S A 100, 6163–6168 (2003).

32. D. A. Dombeck, M. S. Graziano, D. W. Tank, Functional clustering of neurons in motor cortex determined by cellular resolution imaging in awake behaving mice. J Neurosci 29, 13751–13760 (2009).

33. B. Hoeks, Levelt, W.J.M., Pupillary dilation as a measure of attention: A quantitative system analysis. Behavior Research Methods, Instruments, & Computers 25, 16–26 (1993).

34. J. Reimer et al., Pupil fluctuations track fast switching of cortical states during quiet wakefulness. Neuron 84, 355–362 (2014).

35. J. A. Nelder, Wedderburn, W.M., Generalized Linear Models. Journal of the Royal Statistical Society. Series A 135, 370–384 (1972).

36. C. Stringer et al., Spontaneous behaviors drive multidimensional, brainwide activity. Science 364, 255 (2019).

37. S. K. Musall, M. T., Gluf, S., Churchland, A. K., Single-trial neural dynamics are dominated by richly varied movements. biorXiv, https://doi.org/10.1101/308288 (2019).

38. B. Mimica, B. A. Dunn, T. Tombaz, V. Bojja, J. R. Whitlock, Efficient cortical coding of 3D posture in freely behaving rats. Science 362, 584–589 (2018).

39. S. Rozzi, P. F. Ferrari, L. Bonini, G. Rizzolatti, L. Fogassi, Functional organization of inferior parietal lobule convexity in the macaque monkey: electrophysiological characterization of motor, sensory and mirror responses and their correlation with cytoarchitectonic areas. The European journal of neuroscience 28, 1569–1588 (2008).

40. B. A. Olshausen, D. J. Field, Sparse coding of sensory inputs. Curr Opin Neurobiol 14, 481–487 (2004).

41. M. T. Lippert, K. Takagaki, C. Kayser, F. W. Ohl, Asymmetric multisensory interactions of visual and somatosensory responses in a region of the rat parietal cortex. PLoS One 8, e63631 (2013).

42. D. Raposo, M. T. Kaufman, A. K. Churchland, A category-free neural population supports evolving demands during decision-making. Nature neuroscience 17, 1784–1792 (2014).

43. K. Ohki, S. Chung, Y. H. Ch’ng, P. Kara, R. C. Reid, Functional imaging with cellular resolution reveals precise micro-architecture in visual cortex. Nature 433, 597–603 (2005).

44. D. D. Stettler, R. Axel, Representations of odor in the piriform cortex. Neuron 63, 854–864 (2009).

45. E. E. Hecht et al., Differences in neural activation for object-directed grasping in chimpanzees and humans. J Neurosci 33, 14117–14134 (2013).

46. W. Suzuki et al., Mirror Neurons in a New World Monkey, Common Marmoset. Front Neurosci 9, 459 (2015).

47. J. I. Gold, M. N. Shadlen, The neural basis of decision making. Annual review of neuroscience 30, 535–574 (2007).

48. R. A. Andersen, H. Cui, Intention, action planning, and decision making in parietal-frontal circuits. Neuron 63, 568–583 (2009).

49. C. D. Harvey, P. Coen, D. W. Tank, Choice-specific sequences in parietal cortex during a virtual-navigation decision task. Nature 484, 62–68 (2012).

50. B. W. Brunton, M. M. Botvinick, C. D. Brody, Rats and humans can optimally accumulate evidence for decision-making. Science 340, 95–98 (2013).

51. A. S. Morcos, C. D. Harvey, History-dependent variability in population dynamics during evidence accumulation in cortex. Nature neuroscience 19, 1672–1681 (2016).

52. E. J. Hwang, J. E. Dahlen, M. Mukundan, T. Komiyama, History-based action selection bias in posterior parietal cortex. Nat Commun 8, 1242 (2017).

53. A. Akrami, C. D. Kopec, M. E. Diamond, C. D. Brody, Posterior parietal cortex represents sensory history and mediates its effects on behaviour. Nature 554, 368–372 (2018).

54. Q. Wang, O. Sporns, A. Burkhalter, Network analysis of corticocortical connections reveals ventral and dorsal processing streams in mouse visual cortex. J Neurosci 32, 4386–4399 (2012).

55. K. Hovde, M. Gianatti, M. P. Witter, J. R. Whitlock, Architecture and organization of mouse posterior parietal cortex relative to extrastriate areas. The European journal of neuroscience 10.1111/ejn.14280 (2018).

56. R. L. Reep, G. S. Goodwin, J. V. Corwin, Topographic organization in the corticocortical connections of medial agranular cortex in rats. The Journal of comparative neurology 294, 262–280 (1990).

57. R. L. Reep, H. C. Chandler, V. King, J. V. Corwin, Rat posterior parietal cortex: topography of corticocortical and thalamic connections. Experimental brain research 100, 67–84 (1994).

58. S. P. Wise, D. Boussaoud, P. B. Johnson, R. Caminiti, Premotor and parietal cortex: corticocortical connectivity and combinatorial computations. Annual review of neuroscience 20, 25–42 (1997).

59. G. Rizzolatti, L. Fogassi, V. Gallese, Parietal cortex: from sight to action. Curr Opin Neurobiol 7, 562–567 (1997).

60. L. Bonini, P. F. Ferrari, Evolution of mirror systems: a simple mechanism for complex cognitive functions. Ann N Y Acad Sci 1225, 166–175 (2011).

61. L. Fadiga, L. Fogassi, G. Pavesi, G. Rizzolatti, Motor facilitation during action observation: a magnetic stimulation study. J Neurophysiol 73, 2608–2611 (1995).

62. T. F. Roberts, S. M. Gobes, M. Murugan, B. P. Olveczky, R. Mooney, Motor circuits are required to encode a sensory model for imitative learning. Nature neuroscience 15, 1454–1459 (2012).

63. S. M. Wilson, A. P. Saygin, M. I. Sereno, M. Iacoboni, Listening to speech activates motor areas involved in speech production. Nature neuroscience 7, 701–702 (2004).

64. B. Wicker et al., Both of us disgusted in My insula: the common neural basis of seeing and feeling disgust. Neuron 40, 655–664 (2003).

65. T. Singer et al., Empathy for pain involves the affective but not sensory components of pain. Science 303, 1157–1162 (2004).

66. C. Bennett et al., Higher-Order Thalamic Circuits Channel Parallel Streams of Visual Information in Mice. Neuron 102, 477–492 e475 (2019).

67. P. Zhou et al., Efficient and accurate extraction of in vivo calcium signals from microendoscopic video data. Elife 7 (2018).

68. J. Friedrich, P. Zhou, L. Paninski, Fast online deconvolution of calcium imaging data. PLoS Comput Biol 13, e1005423 (2017).

69. L. Sheintuch et al., Tracking the Same Neurons across Multiple Days in Ca(2+) Imaging Data. Cell Rep 21, 1102–1115 (2017).

70. J. C. Dunn, Well-Separated Clusters and Optimal Fuzzy Partitions. Journal of Cybernetics 4, 95–104 (1974).

71. A. Cardona et al., TrakEM2 software for neural circuit reconstruction. PLoS One 7, e38011 (2012).

72. G. Paxinos, Franklin K. (2012) Paxino’s and Franklin’s the Mouse Brain in Stereotaxic Coordinates. (Academic Press).

